# The orientation of cholesterol’s hydroxyl group affects its membrane dynamics and intracellular transport

**DOI:** 10.64898/2026.02.02.703222

**Authors:** Line Lauritsen, Amanda Therese Pauli, Miklas P. W. Larsen, Peter Reinholdt, Holger A. Scheidt, Yuanjian Xu, Douglas F. Covey, Laura Depta, Hogan Bryce-Rogers, Luca Laraia, Jacob Kongsted, Daniel Wüstner

**Affiliations:** Department of Biochemistry and Molecular Biology, University of Southern Denmark, Campusvej 55, DK-5230 Odense M, Denmark; Department of Physics, Chemistry and Pharmacy, University of Southern Denmark, Campusvej 55, DK-5230 Odense M, Denmark; Institute for Medical Physics and Biophysics, Leipzig University, Härtelstr. 16-18, D-04107; Department of Developmental Biology, Washington University, St. Louis, MO 63110, USA; The Taylor Family Institute for Innovative Psychiatric Research, Washington University, St. Louis, MO 63110, USA; Department of Chemistry, Technical University of Denmark, Kemitorvet 207, 2800 Kongens Lyngby, Denmark

**Keywords:** Cholesterol transport, epimers, trafficking, microscopy, fluorescence, probes

## Abstract

The brain, though less than 10% of body mass, contains about 25% of total cholesterol (CHL), emphasizing CHL’s key role in neuronal function. Many CHL actions are stereospecific, as shown by differences from its 3*α*-hydroxy epimer, epicholesterol (epiCHL). How this minor structural change alters membrane properties and sterol transport remains unclear. Here, we compare fluorescent analogs of CHL (cholestatrienol, CTL) and epiCHL (epicholestatrienol, epiCTL), which closely mimic their natural counterparts. Biophysical membrane properties, such as flip-flop, acyl-chain ordering, and interbilayer transfer, depend on the orientation of the 3-hydroxy group. Similarly, transport by sterol transport proteins (STPs) and intracellular trafficking of the sterols in human astrocytes are stereospecific. Treatment with 25-hydroxycholesterol increases uptake of both epimers, but only CTL shows enhanced esterification and lipid droplet storage. These findings demonstrate that subtle cholesterol structural changes affect cellular homeostasis and establish epiCTL as a useful probe of sterol stereospecificity and trafficking.

## Introduction

Cholesterol (CHL) is the most abundant lipid species in the brain, where it is essential for isolating neurons with myelin sheaths and for regulating synaptic activity(1). CHL also controls the activity of membrane receptors and ion channels, and thereby directly impacts cognitive function(2–4). CHL is also the precursor of steroid hormones and oxysterols, which play important roles in a variety of processes, including recovery from injuries and stroke, preventing development of neurodegenerative diseases and maintenance of sterol homeostasis and efflux from the brain(5–9).

Many of the effects of CHL on receptor activity are dependent on specific interactions, which are reduced upon structural changes in the sterol molecule. For example, the orientation of the 3-hydroxy group is important (Fig. 2A), as CHL’s epimer with a 3-hydroxy group in alpha orientation, epicholesterol (epiCHL), often cannot replace CHL in regulating protein activity(10–12). Complicating the matter, CHL also indirectly affects membrane proteins by regulating the biophysical properties of the surrounding lipid bilayer(13– 15). This includes CHL’s well known condensing effect, which results in straightening of fatty acyl chain and thereby tighter packing of phopspholipids in the fluid phase. As a consequence, mechanical properties of the bilayer change, including an alteration of the lateral pressure profile, which in principle can change the conformational dynamics of receptors and thereby their activity(14–16). Interestingly, not only the interaction with receptors but also to some extent the impact on bulk lipid properties differ between CHL and epiCHL. This has for example been shown for the ability to condense membranes, and was assessed by the membrane dipole potential, which is lower for epiCHL compared to CHL, likely due to differences in hydrogen-bonding at the bilayer/water interface(17, 18). Similarly, epiCHL mixes less effectively with saturated phosphatidylcholine in model membranes and therefore has a slightly lower ability to induce the biologically relevant liquid-ordered phase, particularly in vesicles with asymmetric lipid composition between the two leaflets(19, 20).

In contrast to our understanding of the differing effects of CHL and epiCHL on protein and lipid conformations in membranes, very little is known about the role of CHL’s stereochemistry on its trafficking in cells. Why has nature preferred the natural stereochemistry of the 3-hydroxy group of cholesterol? We argue that, besides control of membrane biophysical properties and stereospecific interaction with membrane receptors, the orientation of the 3-hydroxy group could impact sterol trafficking and metabolism. Profiling of epiCHL could help uncover which sterol transport proteins (STPs) and which sterol-mediated processes are actually tolerant of the epimerisation, with implications for e.g., drug design. For comparison, it has been established that the biophysical properties of CHL compared to its more polar derivatives, such as oxysterols, have a large impact on sterol trafficking and metabolism in cells. For example, CHL but not oxysterols, such as 27- or 25-hydroxycholesterol (25HC), resides preferentially in the plasma membrane (PM), while both types of sterols can regulate synthesis and esterification of CHL in the endoplasmic reticulum(21–25). Side-chain oxidized CHL derivatives like 27- or 25HC are in general much more potent than CHL in inducing these effects, likely because they interact more specifically with receptors(26) and due to their differing affinity to STPs(27). Additionally they could be more accessible to STPs due to higher partitioning into the aqueous phase(28, 29). The latter also results in much more rapid intracellular trafficking of these oxysterols compared to CHL(30, 31). The lack of comparable insight into subcellular transport of epiCHL is largely due to a lack of suitable probes to study the trafficking of CHL epimers. Fluorescent analogs of CHL often contain attached dyes, which change the probe’s properties, thereby potentially masking small structural differences. Here, we present a novel intrinsically fluorescent analog of epiCHL based on the well-known cholestatrienol (CTL), which only contains two additional double bonds in the sterol ring system (Fig. 2A). While CTL has the 3-hydroxy group in *β*-orientation like CHL, the new fluorescent epimer, epiCTL, has the 3-hydroxy group in *α*-orientation, like epiCHL. We show that CTL and epiCTL exhibit similar membrane properties as CHL and epiCHL, that STPs can differentiate between the two epimers, and that the intracellular trafficking of CTL and epiCTL is stereo specific.

## Materials and Methods

### Synthesis of epiCTL

EpiCTL was synthesized from 7-dehydrocholesterol as seen in the synthetic scheme as compound **1** (Fig. 1).

**Fig. 1.**
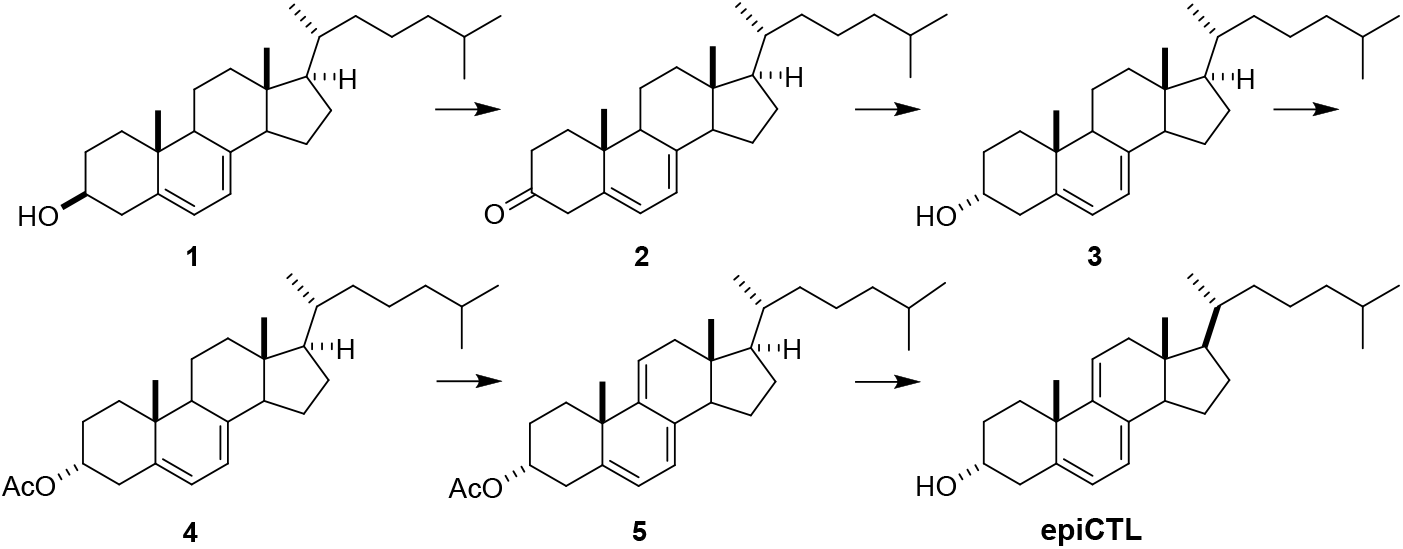
Scheme for syntesis of epiCTL. **1** 7-dehydrocholesterol. **2** Cholesta-5,7-dien-3-one. **3** Cholesta-5,7-dien-3*α*-ol. **4** 3*α*-Acetoxycholesta-5,7-diene. **5** 3*α*-Acetoxycholesta-5,7,9(11)-triene. **epiCTL** Cholesta-5,7,9(11)-trien-3*α*-ol

**Fig. 2.**
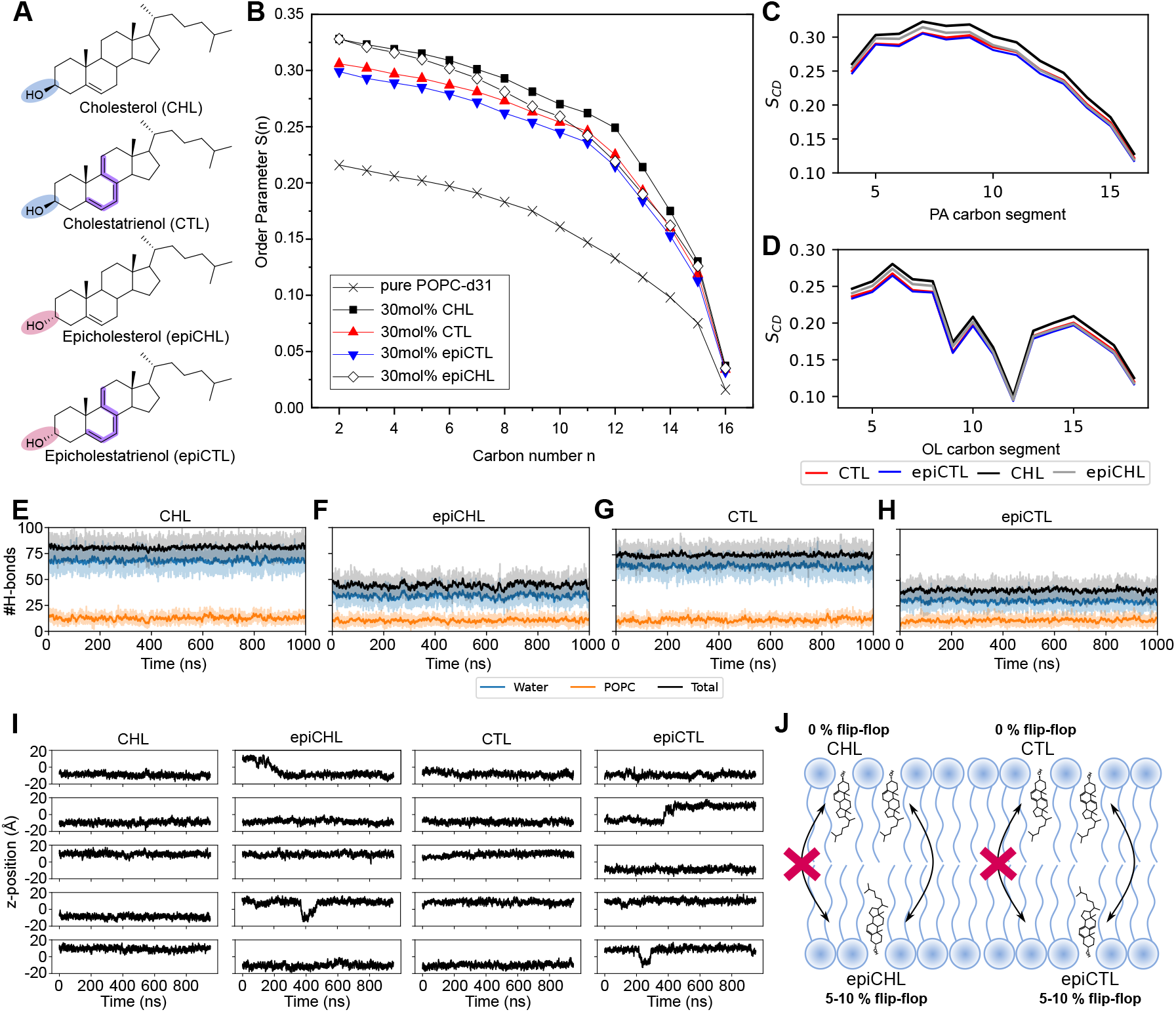
EpiCHL and epiCTL exhibit an increased flip-flop rate in a POPC bilayer compared to CHL and CTL. (A) Chemical structures of CHL, CTL, epiCHL, and epiCTL, where the difference in the position of the 3-hydroxygroup is highlighted with blue and red for the *β*- and *α*-position, respectively. The three conjugated double-bonds making CTL and epiCTL fluorescent are highlighted with purple. (B) ^2^H NMR order parameter S(n) as a function of deuterium position, n, along the POPC acyl chain. (C) Molecular dynamics (MD) simulations of the acyl chain order parameter for the sn-1 chain (palmitic acid) in POPC membranes containing the indicated sterols. (D) Corresponding MD simulations for the sn-2 chain (oleic acid) of POPC (colors correspond to panel C). (E-H) MD simulations of H-bond numbers over time for CHL, epiCHL, CTL, and epiCTL in a POPC bilayers. (I) Representative time traces of the sterol z-position across the bilayer plotted over 1000 ns, for CHL, epiCHL, CTL, and epiCTL. (J) Schematic summary of observed flip-flop behavior, showing higher frequencies for epiCHL and epiCTL.

#### Cholesta-5,7-dien-3-one (2)

To a solution of 7-dehydrocholesterol (**1**, 500 mg, 1.3 mmol) in CH_2_Cl_2_ (20 mL) was added solid NaHCO_3_ (1.5 g) and Dess-Martin periodinane (1.1 g, 2.6 mmol) at room temperature. After 1 h, aqueous NaHCO_3_ (50 mL) was added and the product was extracted into CH_2_Cl_2_ (2 x100 mL). The solvent was removed, and the residue was purified by flash column chromatography (silica gel, eluted with 5-10% EtOAc in hexanes) to give steroid **2** (156 mg, 31%): ^1^H NMR (400 MHz, CDCl_3_) *δ* 5.59-5.57 (m, 1H), 5.40-5.39 (m, 1H), 3.26-3.21 (m, 1H), 2.97-2.92 (m, 1H), 2.48-0.91 (m, 32H), 1.06 (s, 3H), 0.62 (s, 3H); ^13^C NMR (100 MHz, CDCl3) *δ* 211.1, 141.3, 136.2, 120.2, 116.4, 55.9, 54.8, 46.2, 43.3, 42.6, 39.4, 39.1, 37.1, 36.8, 36.1, 36.0, 32.9, 28.1, 28.0, 23.9, 23.0, 22.8, 22.5, 21.6, 18.8, 18.2, 11.9.

#### Cholesta-5,7-dien-3α-ol (3)

To a solution of steroid **2** (156 mg, 0.41 mmol) in THF (20 mL) was added K-selectride (1.0 M in THF, 2 mL, 2 mmol) at -78°C. After 2 h, 3 N NaOH (10 mL) and H_2_O_2_ (2 mL) were added. After addition, the mixture was warmed to room temperature for 1 h. The product was extracted into EtOAc (2 x 100 m). The combined organic layers were washed with brine (3 x 50 mL). The solvent was removed, and the residue was purified by flash column chromatography (silica gel, eluted with 20% EtOAc in hexanes) to give steroid **3** (31 mg, 20%): ^1^H NMR (400 MHz, CDCl3) *δ* 5.63-5.61 (m, 1H), 5.41-5.40 (m, 1H), 4.09 (s, 1H), 2.61-0.92 (m, 35H), 0.95 (s, 3H), 0.63 (s, 3H); ^13^C NMR (100 MHz, CDCl3) *δ* 141.6, 137.3, 121.6, 116.3, 66.2, 55.8, 54.5, 46.2, 42.8, 39.5, 39.2, 38.4, 37.4, 36.1, 36.0, 33.7, 28.9, 28.1, 28.0, 23.8, 23.0, 22.8, 22.5, 20.8, 18.8, 15.9, 11.8.

#### 3α-Acetoxycholesta-5,7-diene (4)

To a solution of steroid **3** (31 mg, 0.081 mmol) in CH_2_Cl_2_ (5 mL) was added acetic anhydride (0.09 mL, 1 mmol) and DMAP (10 mg) at room temperature. After 16 h, the solvent was removed, and the residue was purified by flash column chromatography (silica gel, eluted with 10% EtOAc in hexanes) to give steroid **4** (35 mg, ∼100%): ^1^H NMR (400 MHz, CDCl_3_) *δ* 5.54-5.53 (m, 1H), 5.41-5.40 (m, 1H), 5.06 (s, 1H), 2.55-0.92 (m, 34H), 2.04 (s, 3H), 0.95 (s, 3H), 0.63 (s, 3H); ^13^C NMR (100 MHz, CDCl_3_) *δ* 170.9, 141.3, 137.7, 120.0, 116.4, 69.5, 55.8, 54.6, 45.8, 42.8, 39.5, 39.1, 37.3, 36.1, 36.0, 34.8, 33.9, 28.1, 28.0, 26.0, 23.9, 23.0, 22.8, 22.5, 21.4, 20.7, 18.8, 16.0, 11.8.

#### 3α-Acetoxycholesta-5,7,9(11)-triene (5)

Mercuric acetate (318 mg, 1 mmol) was stirred in HOAc (5 mL) for 15 min. A solution of steroid **4** (35 mg, 0.081 mmol) in CH_2_Cl_2_ (3 mL) was added and the reaction was stirred at room temperature for 16 h. The reaction was added to water and the product extracted into EtOAc (150 mL). The organic layer was washed with aqueous NaHCO3 (3 x 50 mL), the solvent was removed, and the residue was purified by flash column chromatography (silica gel, eluted with 10% EtOAc in hexanes) to give steroid **5** (9 mg, 26%): ^1^H NMR (400 MHz, CDCl_3_) *δ* 5.62-5.61 (m, 1H), 5.51-5.50 (m, 1H), 5.41-5.40 (m, 1H), 5.06 (s, 1H), 2.74-0.87 (m, 31H), 2.02 (s, 3H), 0.88 (s, 3H), 0.58 (s, 3H); ^13^C NMR (100 MHz, CDCl_3_) *δ* 170.8, 144.1, 140.0, 135.4, 121.9, 118.9, 115.6, 77.2, 71.6, 56.4, 50.9, 43.0, 42.1, 39.5, 36.0, 35.9, 35.8, 35.0, 30.2, 29.7, 28.5, 28.0, 26.7, 23.9, 22.8, 22.5, 21.5, 18.4, 11.4.

#### Cholesta-5,7,9(11)-trien-3α-ol (epiCTL)

To a solution of steroid **5** (9 mg, 0.021 mmol) in MeOH (5 mL) was added K_2_CO_3_ (100 mg) at room temperature. The reaction was refluxed for 16 h, MeOH was removed, and the residue was purified by flash column chromatography (silica gel, eluted with 20% EtOAc in hexanes) to give epiCTL (5 mg, 62%): 1H NMR (400 MHz, CDCl_3_) *δ* 5.73-5.72 (m, 1H), 5.52-5.51 (m, 1H), 5.40-5.39 (m, 1H), 4.08 (s, 1H), 2.84-0.92 (m, 32H), 0.88 (s, 3H), 0.57 (s, 3H); ^13^C NMR (100 MHz, CDCl_3_) *δ* 144.4, 140.2, 135.9, 122.2, 120.4, 115.1, 68.6, 56.4, 43.0, 42.1, 39.5, 39.4, 36.0, 35.9, 35.7, 29.7 (2 x C), 29.6, 29.4, 28.5, 28.0, 23.9, 22.8, 22.7, 22.6, 18.4, 11.4.

### NMR spectroscopy

1-palmitoyl-d_31_-2-oleoyl-sn-glycero-3-phosphocholine (POPC-d_31_) and CHL were purchased from Avanti Polar Lipids, Inc. (Alabaster, Al). The desired molar ratio of the respective molecules was first mixed in ethanol. For overnight lyophilization the solvent was evaporated, and the samples were redissolved in cyclohexane. The obtained fluffy powder was hydrated with 50 wt% H_2_O-buffer (10 mM Hepes, 100 mM NaCl, pH 7.4) and equilibrated by ten freeze-thaw cycles. The static ^2^H NMR spectra were acquired on a Bruker (Bruker Biospin, Rheinstetten, Germany) DRX300 NMR spectrometer using the quadrupole echo pulse sequence(32) on a high-power probe with a 5 mm solenoid sample coil. The relaxation delay was 1 s, the delays between the 90° pulses (of ~3.2 µs) were 50 µs. Using the depaked spectra(33), smoothed order parameter profiles were calculated(34). The measurements were carried out at a temperature of 25°C.

#### Molecular dynamics simulations

Membrane bilayers containing in each leaflet 70 POPC lipids and 30 sterol molecules were assembled using packmol. Membranes were assembled for CHL, epiCHL, and their fluorescent analogs. The systems also included 10,000 water molecules, 22 K^+^, and 22 Cl^−^ ions, corresponding to a concentration of about 120 mM KCl. The water molecules were described by the TIP3P force field(35), while the POPC lipids and CHL were described by the amber lipid14 force field(36). Force field parameters for CHL, CTL, epiCHL, and epiCTL were derived using a force matching procedure. An initial parameter set was assigned from GAFF(37), with charges assigned using ESP-fitting, based on r2scan-3c calculations with Orca(38), version 5.0.3. Force-field parameters were then derived using the ForceBalance(39) program, based on quantum-mechanical energies and gradients evaluated using r2scan-3c(40) calculations using Orca. All bonded parameters were optimized, and symmetry-equivalent parameters were assigned equal parameters. We employed an iterative procedure, where the current iteration of the force field was used to run a 10 ns gas-phase stochastic dynamics trajectory with Gromacs(41), from which 100 structures were extracted in each iteration. Quantum-mechanical energies and gradients were evaluated on these structures, and the force-field parameters were then updated to reproduce the complete set of data as well as possible. We repeated this procedure for 10 iterations with a target temperature of 300K in the stochastic dynamics sampling, followed by an additional 5 iterations at an elevated temperature of 1000K, to obtain a better coverage of the parameter space.

All molecular dynamics simulations were carried out using the Gromacs program(41), version 2023.1. The membranes were equilibrated in a three-stage procedure. First, the structures were minimized for 5000 steps with a steepest descent minimizer to remove any bad contacts. Next, we carried out NVT equilibration (200 ps), and NPT equilibration (2 ns) was carried out. A time-step of 2 fs was used. Long-range electrostatics were treated with the particle mesh Ewald method(42) with a cutoff distance of 12 Å. Bonds involving hydrogens were constrained using LINCS(43). The temperature was controlled by a Nose-Hoover thermostat(44, 45) (towards 298.15K), while the pressure (in the NPT simulations only) was controlled by a semi-isotopic Berendsen barostat(46) (towards 1 bar). After equilibration, production molecular dynamics runs (1000 ns) were carried out. In the production simulations, we switched to a Parrinello-Rahman barostat(47). The resulting trajectories were analyzed using Gromacs (for deuterium order parameters), and in-house Python scripts relying on the MDAnalysis library(48) for tilt angle distributions, hydrogen bonding, and autocorrelations of the tilting angle.

### Protein expression constructs

Human ASTER domains of Aster-A(359-547), -B(364-552) and -C(318-504) were subcloned into a pGEX-6p-2rbs vector, thus introducing the cloning artifact ‘GPLGS’(49). The pET22b_His6_STARD1(66-284) and pET22b_His6_STARD3(216-444) plasmids were a gift from James H. Hurley (University of California)(50). The pHIS_2His6_Thrombin_STARD4(2-205;C75S) plasmid was a gift from Young Jun Im (Chonnam National University). STARD5A was a gift from Nicola Burgess-Brown (Addgene plasmid #42392; http://n2t.net/addgene:42392; RRID:Addgene_42392).

### Protein expression and purification

The ASTER domains of human Aster-A(359-547), -B(364-552) and - C(318-504) in pGEX-6p-2rps vectors, including an N-terminal PreScission-cleavable GST-tag were expressed in Escherichia coli (E. coli) OverExpress C41 in Terrific Broth (TB) medium for 16 hrs at 18°C after the induction with 0.1 mM IPTG. Cells were harvested at 3500g for 15 min and lysed by sonication in buffer containing 50 mM HEPES pH 7.5, 300 mM NaCl, 10% (v/v) glycerol, 5 mM DTT, 0.1% (v/v) Triton X-100 and protease inhibitor mix HP plus (Serva). The lysate was purified by affinity chromatography on a GSTrap FF column (Cytiva) using an ÄKTA Start (Cytiva) in buffer containing 50 mM HEPES pH 7.5, 300 mM NaCl, 10% (v/v) glycerol, 5 mM DTT, and 0.01% (v/v) Triton X-100. The GST-tag was cleaved overnight on the column at 4°C. The Aster sterol binding domains were further purified by size-exclusion chromatography (SEC) on a HiLoad 16/600 Superdex 75 pg (Cytiva) in buffer containing 20 mM HEPES pH 7.5, 300 mM NaCl, 10% (v/v) glycerol, and 2 mM DTT.

The START domains of human STARD1(66-284), STARD3(216-444), STARD4(2-205; C75S) and STARD5(6-213) harboring an N-terminal His6-Tag were expressed in E. coli BL21(DE3) in Luria-Bertani Broth (LB) medium for approximately 16 hours at 18°C after induction with 0.15 mM IPTG. Cells were harvested at 3,500g for 15 min and lysed by sonication in buffer containing 50 mM HEPES pH 7.5, 150 mM NaCl, 5% (v/v) glycerol, 5 mM DTT, 0.1% (v/v) Triton X-100, and EDTA-free protease inhibitor cocktail (Sigma-Aldrich). The cleared lysate was purified by affinity chromatography on a Ni-NTA Superflow Cartridge (Qiagen) using an ÄKTA Start (Cytiva) in buffer containing 50 mM HEPES pH 7.5, 150 mM NaCl, 5% (v/v) glycerol, 5 mM DTT. START domains were eluted by using elution buffer containing 50 mM HEPES pH 7.5, 150 mM NaCl, 5% (v/v) glycerol, 5 mM DTT, and 500 mM imidazole. Proteins were further purified by SEC on a HiLoad 16/600 Superdex 75 pg (Cytiva) in buffer containing 20 mM HEPES pH 7.5, 150 mM NaCl, 5% (v/v) glycerol, and 2 mM DTT(27).

### Sterol transfer assay

#### Preparation of Vesicles

A 2 mM stock solution of DHE (Avanti Polar Lipids, #810253), CTL, or epi-CTL in absolute ethanol was prepared. Dansyl-PE (1 mL, 1 mg/mL) in chloroform was obtained from Avanti Polar Lipids (#810333A). A stock solution of DOPC in chloroform (10 mg/mL) had previously been prepared from a 25 mg/mL solution (Avanti Polar Lipids #850375C). The following steps all took place in glass-vials covered in aluminium foil, to keep the light-sensitive lipids in the dark as much as possible: The stock solutions were mixed in a molar ratio of 90/10 DOPC/ligand for the donor vesicles and 97.5/2.5 Dansyl-PE for the acceptor vesicles to a final volume of 1 mL in chloroform. Evaporation of the solvent under a stream of nitrogen, followed by drying under vacuum overnight, afforded the dried lipid films. The lipid films were hydrated in a buffer of 20 mM HEPES pH 7.5, 300 mM NaCl, and 2 mM DTT to a final concentration of 260 µM. The solutions of the lipid films were vortexed extensively until full hydration was observed and sonicated for five min in a 40°C water bath, followed by five freeze-thaw cycles (-196°C to 40°C). Homogeneous unilamellar vesicles were obtained by extrusion 13 times through a polycarbonate membrane (0.1 µm pore size, Avanti Polar Lipids) at 40°C. Solutions were kept on ice and used on the same day as preparation.

#### Microplate-based cholesterol transfer assay

In a non-binding clear-bottom 96-well plate (Greiner Bio-one, cat# 655906), wells were prepared as follows:

### Preparation for run

A master mix of donor and acceptor liposomes was made in buffer (20 mM HEPES pH 7.5, 300 mM NaCl, 2 mM DTT) affording a final assay concentration for both donor and acceptor as 12.5 µM (total liposome concentration 25 µM) and a final assay volume of 100 µL. For compound containing runs, protein and compound were incubated at a concentration 20 times the desired assay concentration for 15 min.

### The run

95 µL of liposome mixture was then added to each well (maximally 6 wells per run). Fluorescence intensity measurements were performed in a Tecan Spark Cyto plate reader at 25°C, measuring from the bottom at 10 sec intervals. The excitation filter was set at 340 *±* 20 nm, and the emission filter was set to 535 ± 20 nm. After approximately 2 minutes, the measurement was paused, the plate was ejected, and 5 µL protein (or protein + compound pre-mix) was added as quickly as possible and mixed with a pipette to a desired final concentration. The measurement was continued, and the total measuring time was 12 min. Data was normalised to I_0_ of the donor + acceptor (before adding protein). Where spurious fluorescence quenching was observed upon protein addition, data was normalized to the first timepoint after addition (t = 0.25 min). All data was plotted in GraphPad Prism 5.

### Cell line

Immortalized human astrocytes (IM-NHA, # P10251-IM, Innoprot) were grown at 37°C, 5% CO_2_, and 100% humidity, and in astrocyte medium (AM, # P60101, Innoprot) supplemented with 2% Fetal Bovine Serum (FBS), astrocyte growth supplement, and 1% penicillin/streptomycin. Media was changed every second day and subcultured at 90% confluence using 1X trypsin-EDTA (# T47174, Sigma-Aldrich). Cells were kept in M1 media (150 mM NaCl (Merck, # 1.06404.1000), 5 mM KCl (Merck, # 104936), 1 mM CaCl_2_ (Merck, # 2382.1000), 1 mM MgCl_2_ (Merck, # 1.05833.1000), 5 mM glucose (Merck, # 1.08342.1000), and 20 mM HEPES (Sigma, # H3375-100G), pH addjusted to 7.4) under imaging.

#### Labeling cells with CTL and epiCTL

IM-NHA cells were plated in 35mm microscope dishes (MatTek, # TKO-P351184-408) for sterol trafficking measurements the day prior to loading with the fluorescent sterols. CTL and epiCTL were loaded onto fatty-acid free bovine serum albumin (BSA, Sigma-Aldrich, # A7030-50G). In short this was done by adding 2 · 10−7 mol of the sterol to a clean glass vial, evaporating the solvent before resuspending it in 10 µL of absolute ethanol and then adding 990 µL of 50 mg/mL BSA in M1-media, resulting in a final concentration of 200 µM of the sterol. Then the solution was vortexed for 5 min, before it was allowed to equilibrate for 0.5-1 hour at room temperature in the dark. The final concentration of ethanol was under 1 % (v/v). For trafficking measurements 24 hrs of uptake IM-NHA were incubated with 20 µM of the sterol/BSA complex and ∼ 25µg/mL Rhodamine Dextran (RhDex, Invitrogen™, # D1819) in LPDS media, DMEM high glucose (Sigma-Aldrich, # D6429-500ML) containing 10% lipoprotein-deficient serum (LPDS, Merck, # S5519-50ML) and 68.8 nM Na-oleate (Sigma-Aldrich, # O7501). For treatment with avasimibe (Bionordika, # CC-18129-10) and/or 25HC (Sigma-Aldrich, # H1015) they were both added in a concentration of 20 µM together with the addition of sterol/BSA. The day of imaging the media was exchanged to M1 media, and 10 min incubation at 37°C with 1µM Bodipy™ 493/503 (Thermo Fisher Scientific, # D3922) was used to stain the LDs. The IM-NHA cells were then washed three times before imaging. For the cells prepared for 24 hrs efflux measurements were loaded with 20 µM sterol/BSA in LPDS media one day after plating. 24 hrs after loading the media was exchanged to fresh LPDS media, and ∼ 25µg/mL RhDex was used for stainng LE/LYs, and treatment with avasimibe and/or 25HC was continued. The cells were left to efflux for 24 hrs.

### Sterol live cell imaging

Before imaging, the cells were washed carefully with M1-medium, and the lipid droplets (LDs) were labeled by incubating the cells for 10 min at 37 °C with 1 µM Bodipy (Invitrogen™, # D3922), and then they were washed 3*×* with M1-media. When imaging the endocytic recycling compartment (ERC) Transferrin-CF640R (biotium, # 00085) was used. Here, the cells were incubated at 37 °C in M1-media containing 20 µg/mL Transferrin-CF640R. Bodipy (1 µM) was added after 10 min of incubation and maintained for the final 10 min, followed by 3*×*flushing with M1-media. The cells were imaged at room temperature in M1-media at an UV-optimized widefield epifluorescence Leica DMIRBE microscope using a 63*×* 1.3 NA oil immersion objective (Leica Lasertechnik GmbH) containing a 10*×* extra magnification lens in the emission light path, resulting in a final pixel size of 193 nm. For illumination control, a Lambda SC smart shutter (Sutter Instrument Company) was used. For capturing the images, an Andor Ixon blue EMCCD camera operated at -75°C was driven by the Andor Solis: X-5771 (Solis version: 4.16.30003.0). Transferrin-CF640R signal was captured using the Y5 Leica filter cube with a bandpass filter 620/60 nm as excitation filter, a 660 nm dichromatic mirror, and a 700/75 nm band-pass emission filter. Rh-Dex using the Rhod Et Leica filter cube bandpass filter 530/30 nm as excitation filter, a 560 nm dichromatic mirror, and a 550 nm longpass emission filter. Bodipy was imaged using an L5 Leica filter cube with a band-pass filter 480/40 nm as excitation filter, a 505 nm dichromatic mirror, and a 527/30 nm bandpass emission filter. The sterols CTL and epiCTL were imaged using a custom filter cube from Chroma Technology with a bandpass 335/20 nm excitation filter (Chroma, # D335/20x), a 365 nm dichromatic mirror (Chroma, # 365dclp), and a bandpass 405/40 nm emission filter (Chroma, # D405/40m). When imaging the sterols, a 100-frame bleaching stack with an exposure of 0.4 s/frame was recorded, which was used for background and autofluorescence correction, as described below.

### Image analysis of sterol trafficking

All data analysis were carried out using custom-written scripts in MATLAB2023b (The MathWorks, Inc). All UV images of CTL and epiCTL were corrected for autofluorescence by subtracting the last frame in the bleaching stack from the first frame(51, 52), resulting in only the true signal from the probe. To obtain single cell data, the Bodipy signal of the cells was segmented using the built-in cytoplasm 2.0 model (‘cyto2’) in Cellpose(53), which can be run as a function in MATLAB. Here, the model parameters were updated so that they gave the best possible segmentation with a cell diameter of 150, a flow error threshold of 2.5, and a cell threshold of −5. To exclude cell debris, abnormally small cells, and segmentation artifacts, a size threshold was applied such that objects occupying less than 1% of the image were discarded. When staining the cells with organelle-specific markers there will be some background staining of the dyes in the cytosol. Since the cells have a varying amount of dye in the cytosol and thickness, it is favorable to do a local background subtraction to enhance the structures of the organelle. Here, this is done by morphological operations, first an erosion and then a dilation (carried out by the MATLAB function ‘imopen’) with a disk size larger than the organelle, which results in an image of the local background. The local background can then be subtracted from the original image, leaving the structures of the wanted organelles. To detect these in an automated manner, the mean signal of the image is calculated, and the mask is defined as the signal above that mean multiplied by a threshold factor. Since the organelles vary a bit in size and brightness, a disk size of ∼ 1µm and a threshold of 5 for LDs, a disk size of ∼ 2.9µm and a threshold of 2 for RhDex, and a disk size of ∼ 5.8µm and a threshold of 2 for the ERC were used. The masks of the single cells were used to calculate the mean UV intensity per cell. The fraction UV for each organelle was calculated by applying the organelle mask within the single cell, and calculating the mean intensity found in the organelle mask divided by the mean of the remaining UV intensity in the cell. Here, a value >1 indicates a higher level of sterol in that organelle, and a value <1 indicates less sterol in that organelle.

## Results

### The orientation of the 3-hydroxy group of CHL affects its membrane properties

To investigate how the orientation of the 3-hydroxy group of CHL and epiCHL and their fluorescent analogs CTL and epiCTL (Fig. 2A) affects the membrane properties, we first examined the impact of these sterols on a POPC bilayer. ^2^H NMR experiments, in which deuterium atoms were substituted along the palmitoyl chain of POPC, were used to determine order parameter in vesicles composed of pure POPC-*d*_31_ or POPC-*d*_31_ containing 30 mol% of CHL, CTL, epiCHL, or epiCTL (Fig. 2B). All four sterols increased the order parameter of the sn1-chain relative to pure POPC-*d*_31_. CHL produced the strongest ordering effect, whereas epiCHL resulted in a slightly reduced order parameter, particularly in the central region of the bilayer. Incorporation of the fluorescent analogs CTL and epiCTL further decreased the order parameter, following a trend similar to CHL and epiCHL, with CTL inducing higher order than epiCTL.

MD simulations supported the NMR findings (Fig. 2C and D). The calculated order parameters for the *sn*-1 (palmitric acid, PA) and *sn*-2 (oleic acid) chains again showed that CHL induced the highest order, followed by epiCHL, with CTL and epiCTL only marginally lower. Analysis of hydrogen bonding revealed that all sterols formed a comparable number of hydrogen bonds (H-bonds) to the POPC lipids (Fig. 2E-H). However, CHL and CTL established significantly more H-bonds to water compared to their epimers, resulting in an overall reduction in the total number of H-bonds for epiCHL and epiCTL to nearly half of CHL and CTL.

Next we analyzed the sterol position within the bilayer over 1000-ns long MD simulation. On this timescale CHL and CTL remained stably in one side of the bilayer without signs of flip-flop (Fig. 2I). In contrast, both epiCHL and epiCTL exhibited flip-flop events, which are visible as abrupt changes in their z-position (Fig. 2I). We define a change in z-position as flip-flop event, if the sterol has been moved from z > 10Åto z < -10Åacross the bilayer. A summary of flip-flop frequencies are shown in Fig. 2J, which reveals that neither CHL nor CTL migrate to the opposite leaflet during the MD simulations. In contrast, both epiCHL and epiCTL underwent flip-flop events in 7.7 ± 2.1% of the traces, and epiCHL in 6.6 ± 1.4% of the traces. These are calculated based on 3 individual simulation runs of 60 traces. The increased flexibility of epiCHL and epiCTL is likely due to the reduced number of H-bonds formed in the membrane, resulting in a less constrained bilayer positioning.

### Passive and protein-mediated sterol transfer between vesicles is stereospecific

To examine how the orientation of the 3-hydroxy group influences protein-mediated sterol transfer, we employed an *in vitro* FRET assay. Donor liposomes (L_D_) containing CTL or epiCTL served as FRET donors, while acceptor liposomes (L_A_) were loaded with Dansyl-PE. Mixing L_D_ and L_A_ liposomes allowed monitoring of the fluorescence ratio I/I_0_ over time. At t=0, a sterol binding domain of a STP was added (Fig. 3A). An increase in I/I_0_ indicated transfer of the fluorescent sterol from L_D_ to L_A_, whereas a constant signal reflected no transfer.

**Fig. 3.**
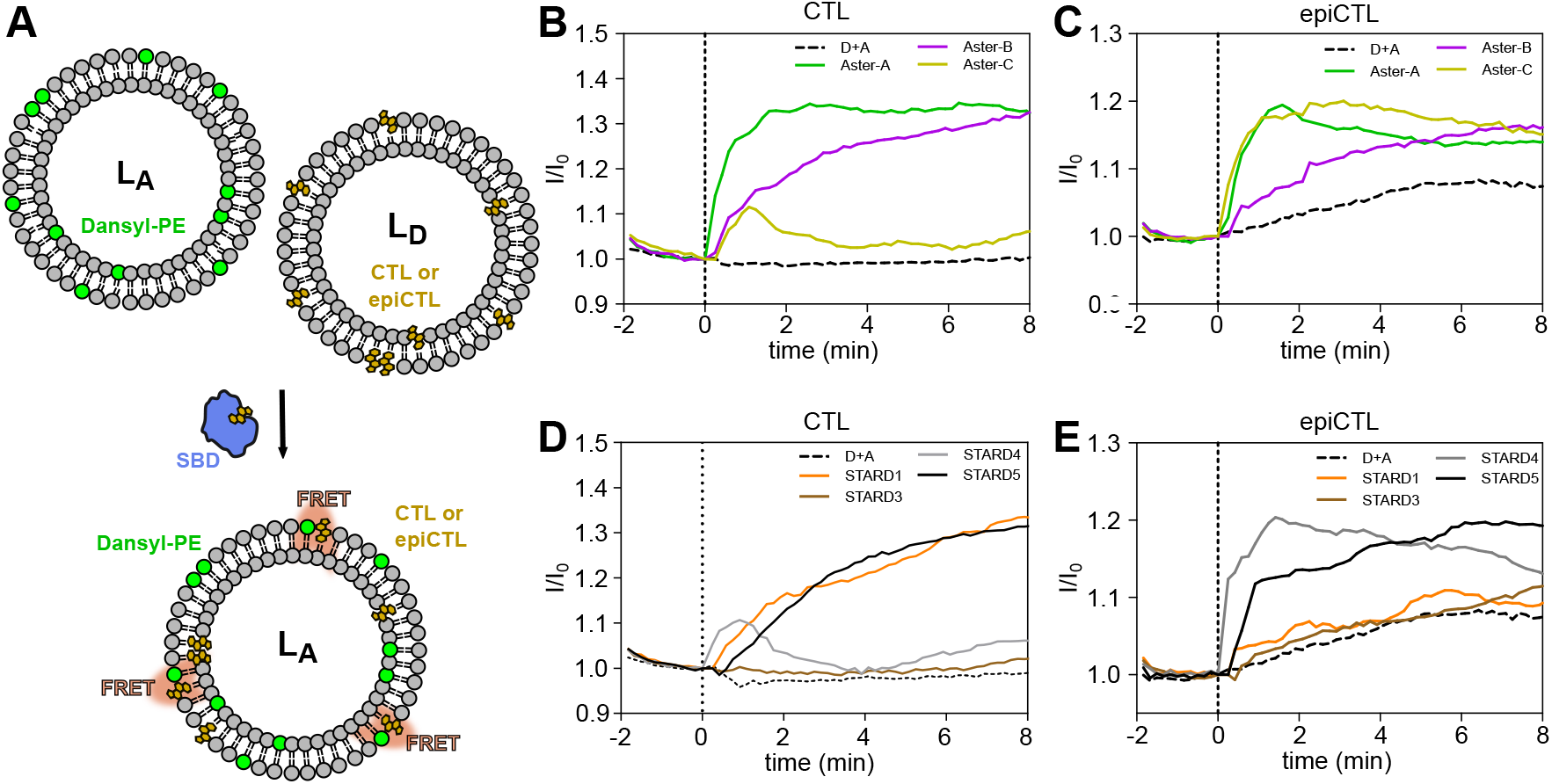
Stereospecific transport of CTL and epiCTL by sterol-binding domains (SBDs). (A) Schematic of the FRET-based liposome transport assay. Donor liposomes (L_D_) containing CTL or epiCTL were mixed with Dansyl-PE-loaded acceptor liposome (L_A_), and transfer was monitored by changes in I/I_0_. (B,C) Transport of CTL (B) and epiCTL (C) by 1 µM Aster-A/B/C. (D,E) Transport of CTL (D) and epiCTL (E) by 1 µM STARD1/3/4/5. Data shown are a representative from two biological replicates (n = 2).

Two classes of STPs were investigated(54). The first class comprised of the Aster proteins, which facilitate CHL transport from the plasma membrane to the endoplasmic reticulum(55). CTL was transferred most efficiently by Aster-A, less efficiently by Aster-B, and only partially by Aster-C (Fig. 3B). In contrast, epiCTL was transported most efficiently by Aster-A and Aster-C, with Aster-B showing moderate activity (Fig. 3C). The preference of Aster-C for epiCTL is notable as it is in line with its preference for the enantiomer of its sterol-inspired inhibitor astercin-2(56).

The second class of STPs tested was the steroidogenic acute regulatory protein-related lipid transfer (START) family, which is known to mediate intracellular CHL transport(57). CTL was transferred efficiently by STARD1 and STARD5, less efficiently by STARD4, and not at all by STARD3 (Fig. 3D). STARD5 has been reported to bind cholesterol or cholic acid, with recent data suggesting that cholic acid derivatives may be the endogenous ligands. However, these results support the notion that both classes of lipids could be substrates(27, 58). EpiCTL was transported efficiently by STARD5, moderately by STARD4, and not detectable by STARD1 and STARD3 (Fig. 3E). Transport efficiencies of CTL and epiCTL by the eight tested proteins are summarized in SI in Table 1, where they are compared to those of DHE (Fig. S6).

Interestingly, control experiments performed without addition of STPs (dotted line (D+A) in Fig. 3B-E) showed stable I/I_0_ values when the L_D_ liposomes contained CTL (Fig. 3B,D). In contrast, L_D_ liposomes containing epiCTL displayed a consistent increase in I/I_0_ (Fig. 3C,E). This important result suggests that epiCTL has a higher rate of spontaneous sterol exchange between liposomes compared to CTL. Supporting that notion are earlier findings of enhanced inter-bilayer transfer of epiCHL compared to CHL(59). Like the higher flip-flop rate observed for epiCTL (Fig. 2) increased passive inter-membrane exchange is likely a result of less stable membrane interactions of epiCTL compared to CTL. Alternatively, epiCTL could faciliate vesicle fusion, resulting in an apparent raise of sterol transfer between the liposomes.

### Stereospecific distribution of CTL and epiCTL in IM-NHA

Given the stereospecificity observed in protein-mediated sterol transfer, we next examined whether CTL and epiCTL distribute differently within cells. IM-NHA cells were loaded with CTL/BSA or epiCTL/BSA for 24 hrs, followed by either direct imaging (uptake) or a subsequent 24 hrs efflux period. Cells were co-stained for lipid droplets (LD, Bodipy) (Fig. 4A), late endosomes/lysosomes (LE/LYs, RhDex) (Fig. 4A), or the endocytic recycling compartment (ERC, transferrin-CF640R) (Fig. S3). CTL fluorescence intensity was consistently higher than that of epiCTL (Fig. 4A). Quantification of mean fluorescence intensity per cell revealed a nearly five fold higher median intensity for CTL compared to epiCTL (Fig. 4B,C). Analysis of sterol efflux revealed that CTL-treated cells retained ∼ 72% of their initial fluorescence, compared to only ∼ 32% for epiCTL-treated cells (Fig. 4B,C).

**Fig. 4.**
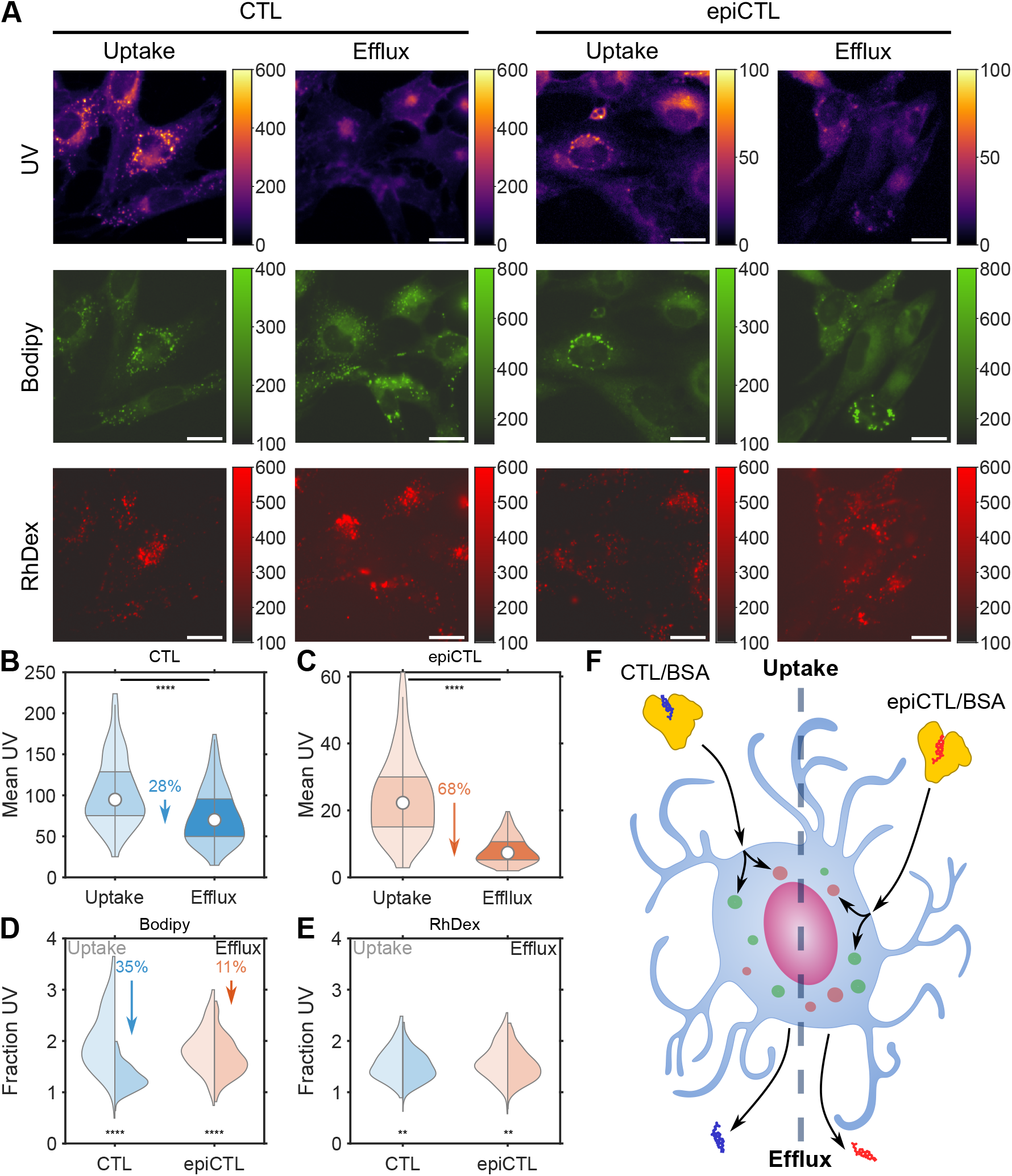
Intracellular distribution of CTL and epiCTL in IM-NHA. (A) Widefield fluorescence images acquired in the UV channel (CTL and epiCTL), green channel (LDs, Bodipy), and red channel (LE/LYs, RhDex) of IM-NHA after 24 hrs uptake and following 24 hrs efflux. Scale bars, 20 µm. (B) Violin plot of mean UV intensity per cell for CTL uptake (*N*_*cells*_ = 277) and efflux (*N*_*cells*_ = 407). (C) Violin plot of mean UV intensity per cell for epiCTL uptake (*N*_*cells*_ = 310) and efflux (*N*_*cells*_ = 333). (D) Double-sided violin plot showing the fraction of UV signal localized in the LDs during uptake (left) and efflux (right) for CTL and epiCTL. (E) Double-sided violin plot showing the fraction of UV signal found in the LE/LYs during uptake (left) and efflux (right) for CTL and epiCTL. (F) Schematic summary of the setup to investigate intracellular distribution for CTL and epiCTL.

To validate that cellular fluorescence reflected comparable amounts of sterol, emission spectra of CTL and epiCTL were recorded at equal concentrations (Fig. S2A,B). Integrated intensities at the start condition were similar, with epiCTL exhibiting slightly higher values (Fig. S2C). Comparable integrated intensities were also obtained for uptake and efflux conditions. When calculating uptake directly from media concentrations, no significant difference between CTL and epiCTL was found (Fig. S2D). This result contrasts with the lower intracellular epiCTL fluorescence observed in imaging (Fig. 4A-C). Similarly, efflux measurements based on spectra suggested lower release of epiCTL compared to CTL, whereas cell-based fluorescence indicated the opposite trend (Fig. S2D).

This discrepancy suggests that epiCTL may undergo intra-cellular conversion to a non-fluorescent derivative. Such a process would lead to an underestimation of epiCTL uptake by imaging and could explain the apparent higher efflux measured in cells. To test whether epiCTL or eventual metabolites derived from epiCTL can affect cellular physiology, intracellular calcium levels were monitored using the calcium sensitive dye, Cal520, as an indicator. Cells treated with epiCTL/BSA displayed a significant elevated calcium concentration compared to cells treated with CTL/BSA and CHL/BSA (Fig. S4). Cells treated with epiCHL/BSA also showed a significant increase in calcium levels compared to treatments with CTL/BSA and CHL/BSA, however not to the same extent as the cells treated with epiCTL/BSA(Fig. S4). These results indicate that epiCTL or its metabolites trigger a specific cellular response.

During uptake a larger fraction of CTL is found in LDs compared to epiCTL (Fig. 4D). Upon efflux, CTL partitioning in LDs decreased with ∼ 35%, whereas epiCTL decreased only ∼ 11% (Fig. 4D). In contrast, partitioning into LE/LYs was similar for both sterols and remained largely unchanged between uptake and efflux (∼ 4% variation in the mean for CTL and ∼ 6% for epiCTL, Fig. 4E). The experimental setup is illustrated in Fig. 4F, showing uptake from BSA, intracellular distribution, and efflux.

#### 25HC selectively promotes esterification and LD accumulation of CTL, but not epiCTL

To probe mechanistic differences in intracellular trafficking, IM-NHA cells were treated with 25HC, which is known to stimulate CHL esterification(60– 62). Treatment with 25HC increased uptake of both CTL and epiCTL (Fig. 5A,B). In contrast, efflux was reduced by ∼ 10% for CTL, whereas epiCTL efflux remained unchanged (Fig. 5A,B, Fig. S5A). Consistent with increased esterification, CTL showed higher accumulation in LDs in cells treated with the oxysterol (Fig. 5C). Notably, 25HC did not alter the fractional distribution of epiCTL in LDs (Fig. 5C).

**Fig. 5.**
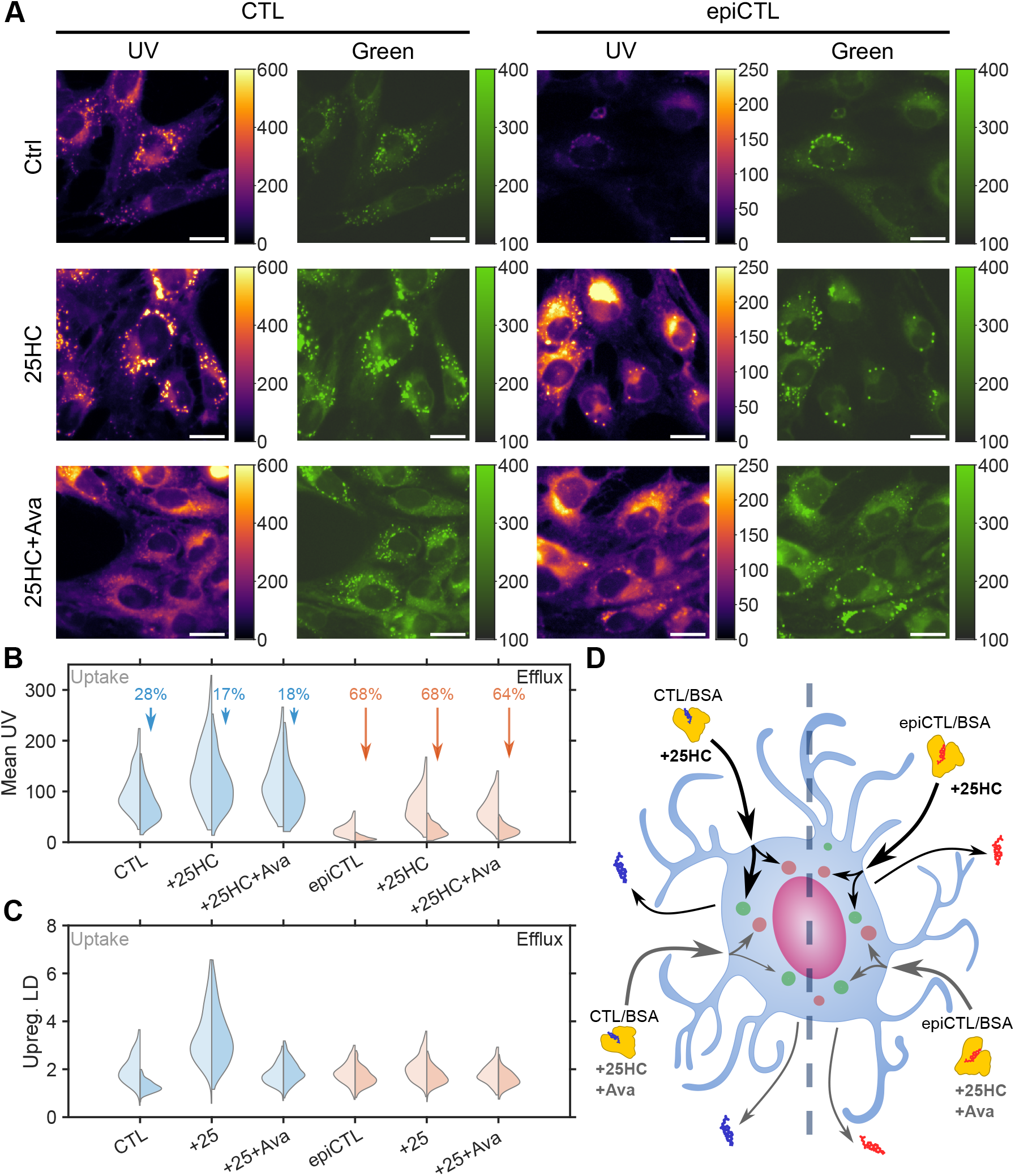
25HC reveals stereospecific intracellular esterification of CTL, but not epiCTL. (A) Widefield fluorescence images of IM-NHA loaded with CTL/BSA or epiCTL/BSA for 24 hrs (uptake) in presence of either sterol alone (ctrl), sterol + 20 µM 25HC (25HC), or sterol + 20 µM 25HC + 20 µM avasimibe (25HC+Ava). Images were acquired in the UV channel (CTL and epiCTL), green channel (LDs, Bodipy). Scale bars are 20 *µ*m. (B) Double sided violin plot showing the distribution of mean UV intensity per cell for uptake (left) and efflux (right) of the conditions shown in A. (C) Double sided violin plot showing the distribution of fraction of UV localized in LDs per cell. Data are from three biological replicates. The number of analyzed cells for each condition is given as (uptake/efflux): CTL, sterol alone (277/407), +25HC (283/396), +25HC+avasimibe (296/402); epiCTL, sterol alone (310/333), +25HC (325/324), +25HC+avasimibe (297/334). (D) Schematic representation of the main differences between CTL and epiCTL trafficking when adding 25HC and avasimibe.

To test whether these effects depend on sterol esterification, cells were treated with avasimibe, an inhibitor of acyl-coenzyme A-acyl transferase (ACAT)(63). Avasimibe did not alter the increased uptake of CTL or epiCTL (Fig. S5B), indicating that the uptake effect is not ACAT-dependent. However, avasimibe reduced the enrichment of CTL in LDs, likely because it blocks the esterification (Fig. 5C). Upon ACAT inhibition, CTL partitioning into LDs became similar to that of epiCTL, suggesting stereospecificity of ACAT, which efficiently esterifies CTL but not epiCTL. These results are in line *in vitro* observations(64), which showed stereospecific activity and allosteric regulation of ACAT by sterols. Fig. 5D shows a schematic summary of our results obtained in cells. Using intrinsically fluorescent analogs of CHL and epiCHL, we find that 25HC stimulates sterol uptake, with no apparent impact on sterol efflux from astrocytes. Enhanced trafficking of cholesterol from the PM to the ER upon treatment with 25HC has been shown previously(65), and we show that this effect is not stereospecific. However, the subsequent sterol esterification by ACAT depends on the orientation of the 3-hydroxy group. Indeed, 25HC increased the localization of CTL in LDs and this could be blocked by inhibiting ACAT using avasimibe, but it didn’t affect the overall uptake or efflux.

## Discussion

In this study, we present epiCTL, a novel fluorescent analog of epiCHL. Using NMR and MD simulations, we compared its effect on membrane properties with CTL, a well established analog of CHL, but also with the natural sterols, CHL and epiCHL. CHL and CTL increased bilayer order parameters and acyl chain order more strongly than their epimers (2B-D). In contrast, epiCHL and epiCTL exhibited higher membrane mobility, reflected in increased flip-flop rates (Fig. 2I,J). We attribute this to the reduced number of hydrogen bonds formed by the epimers, epiCHL and epiCTL, within the bilayer (Fig. 2E-H). This agrees with earlier findings by Róg et al. (66), who demonstrated that a reduced amount of hydrogen bonds at the membrane/water interface correlated with an increase in flip-flop rates comparing CHL and keto-CHL.

Due to the small and compact size of CHL, the addition of fluorophores often alters its biophysical properties(67, 68) and intracellular localization(69, 70). Fluorophore placement and orientation can strongly affect the recognition of CHL by STPs(27). We also demonstrate that CTL and epiCTL mimic the bilayer positioning of CHL and epiCHL (Fig. 2 and S1). The orientation of the 3-hydroxyl group alone dictates the binding and transport by certain STPs. Strikingly, we found that epiCTL has a much higher passive inter-membrane transfer than CTL (Fig. 3), likely due to the less efficient anchoring of epiCTL in the bilayer. Not only the membrane properties but also the recognition and transfer of sterols by STPs are stereospecific. Indeed, Aster C, StARD1, and StARD4 displayed the highest stereospecificity of the tested STPs (Tab. 1), emphasizing that CTL and epiCTL are powerful probes for resolving subtle stereospecific differences in sterol trafficking.

We also observed stereospecific behavior of CTL and epiCTL at the cellular level. Although CTL and epiCTL showed a comparable uptake from media (Fig. S2), CTL fluorescence was approximately fivefold higher per cell (Fig. 4A-C). In parallel, epiCTL - but not CTL - triggered elevated cytosolic calcium levels (Fig. S4). These findings suggest, that epiCTL triggers specific cellular responses which are not activated by either CTL, CHL, and to a lesser extent epiCHL. Also, we cannot rule out that epiCTL is chemically converted in the cells, for example into a non-fluorescent sterol species with particular activity. Subcellular localization of the two probes was largely comparable, with only modest differences in LD partitioning (Fig. 4D).

Our data further suggest that sterol esterification is stereospecific. Treatment with 25HC, a known agonist for ACAT mediated esterification(60–62), enhanced uptake of both CTL and epiCTL, but only promoted LD accumulation of CTL (Fig. 5A,B). This effect was blocked by the ACAT inhibitor avasimibe, confirming that esterification of CTL - but not epiCTL - is ACAT dependent. Interestingly, 25HC did not increase efflux of either sterol under our conditions (Fig. 5B), in contrast to its reported role in enhancing cholesterol efflux in fibroblasts and macrophages via LXR*α*-driven ABCA1 upregulation(71). Future studies should be directed towards exploring stereospecific interaction of sterols with LXRs in living cells, and epiCTL will be an important tool for such experiments.

## Conclusion

Our study reveals that a single structural change of CHL affects its membrane dynamics (number of hydrogen bonds to water and flip flop rate), protein mediated transport, and intracellular transport. We were able to prove that CTL and epiCTL, our intrinsically fluorescent analogs of CHL and epiCHL, were able to mimic the membrane properties of CHL and epiCHL. This allowed us to resolve differences in protein mediated transport between model membranes, revealing that STPs can discriminate between the two epimers. The finding that Aster-C and STARD4 preferentially transport epiCTL over CTL suggests that even a single inversion at the 3-hydroxy position can reshape sterol flux. This stereochemical sensitivity offers new opportunities for designing tailored steroidal ligands that selectively modulate cholesterol transport and metabolism.

Fluorescence microscopy of live astrocytes showed differences in metabolism of the two epimers. CTL was able to be esterified, and this effect was amplified by treatment with 25HC, and diminished by the addition of avasimibe. Whereas epiCTL was metabolized through another pathway converting it to something non-fluorescent that triggered an amplified calcium signal. These findings highlight that stereospecificity extents beyond receptor specificity to core aspects of intracellular trafficking and handling of CHL. Beyond providing fundamental insights, epiCTL proved to be a powerful tool to investigate sterol stereospecificity in living cells.

## ACKNOWLEDGEMENTS

The work presented here is supported by the Carlsberg Foundation, grant CF23-1086. The Wüstner group was further supported by funding from Lundbeck Foundation (grant no. R366-2021-226). We would further like to acknowledge the imaging facility DaMBIC for providing microscopy infrastructure (Novo Nordisk Foundation, grant no. NNF18SA0032928). The Laraia Laboratory was supported by funding from the Novo Nordisk Foundation (grant no. NNF19OC0055818, NNF19OC0058183, NNF21OC0067188), the Carlsberg Foundation (grant no. CF19-0072) and the European Union (ERC, ChemBioChol, 101041783). We thank Mjaftime Ismaili and David Frej Nielsen for their support with the expression and purification of proteins. DFC was supported by NIH grant 1 P50 MH122379.

