## Supplementary Note 1 for "The orientation of cholesterol’s hydroxyl group affects its membrane dynamics and intracellular transport"

### Supplementary Note 1: The orientation of cholesterol's hydroxyl group affects its membrane dynamics and intracellular transport

#### Supplementary methods.

***Spectral measurements of media containing CTL or epiCTL.*** Spectra of 100 µL media containing 20 µM CTL/BSA, epiCTL/BSA, or just BSA diluted with 400 µL of 1X PBS were acquired for on a ISS K2 spectrofluorometer. Here, excitation of 335 nm was used and emission spectra were recorded 360-450 nm, with emission slits of 1.0 mm, time base of 1 s, and a step size of 1 nm. Spectra were recorded from the starting media before addition to the cells (start), from the media after 24 hrs of uptake (uptake), and from the media after efflux (efflux).

***Determination of apparent differences in calcium signaling.*** **Cell preparation:** IM-NHA were plated in 8-well Ibidi chambers (#80807, Ibidi GMBH) for calcium imaging the day prior loading with the sterols. The cells were loaded with 20µM sterol/BSA 24 hrs prior imaging. CHL/BSA and epiCHL/BSA were prepared similar to CTL/BSA and epiCTL/BSA. Labelling with the calcium sensor Cal520-AM (#21130, ATT Bioquest) was carried out right before imaging. Here, the cells were stained with 0.5µM Cal520-AM in M1 media for 60 min at 37°C, and then washed 3 times before imaging.
**Imaging of Cal520:** Images of our calcium sensor, Cal520, were recorded on a Ti2-Nikon widefield microscope using 488 nm excitation from a LED-illumination source (CoolLED pE-300) through a FITC filter cube (excitation filter: 467-498 nm, dichroic mirror: 506 nm, emission filter: 513-556 nm). Images were captured using a Plan Apo $\lambda$ 20X objective with a NA of 0.75 and 80 ms of exposure on an Andor Zyla VSC-00580 camera resulting in a pixel size of 323.6 nm.
**Calcium analysis:** Analysis of the Cal520 intensities was carried out in MATLAB, where the cells were segmented using the pretrained model cyto2 in Cellpose, with model parameters being a cell diameter of 100, flow error threshold of 1, cell threshold of -2, and normalized images. From this mask the median Cal520 signal per cell could be calculated and apparent calcium levels for the four different conditions can be visualized in Fig. S4E.

#### MD simulations.

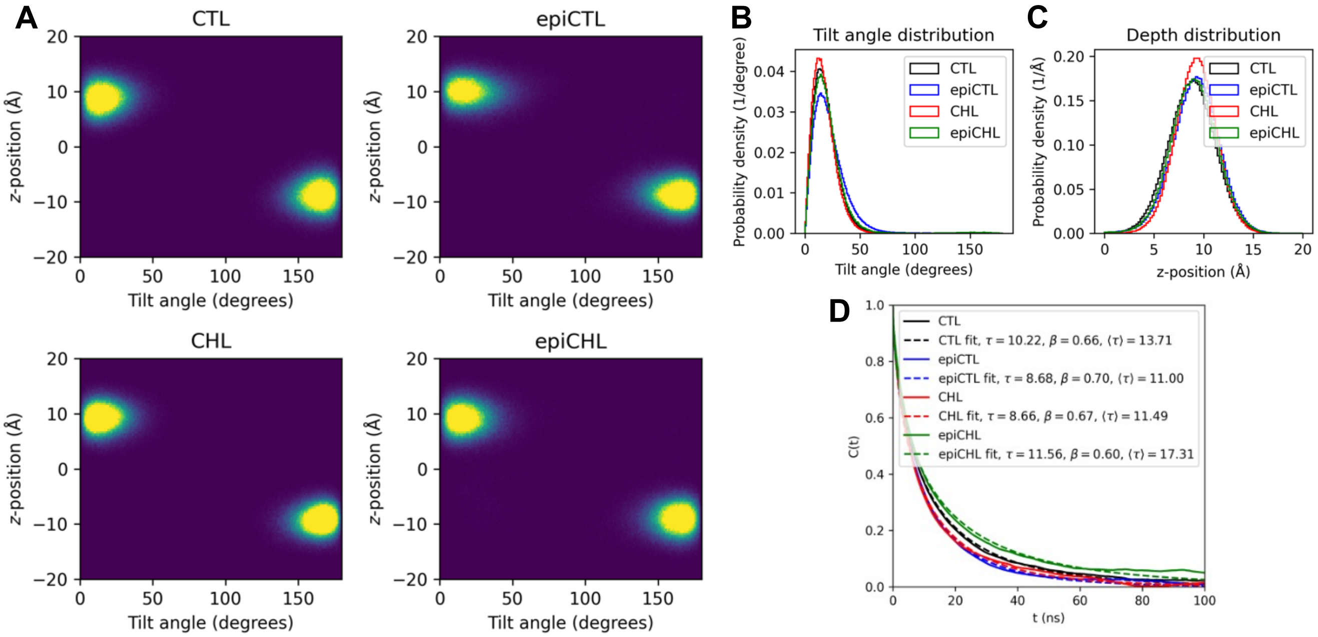


**Fig. S1.** Sterol organization and dynamics in lipid membranes. (A) 2D density heat maps of CTL, epiCTL, CHL, and epiCHL showing their distance from the center of the membrane (Å) plotted against the tilt angle (degrees). (B) Tilt angle distributions of the four sterols. (C) Bilayer depth distributions of the four sterols. (D) Autocorrelation of the tilting angle plotted over time.

**Spectra of CTL and epiCTL conjugated on BSA.** To address whether the uptake and efflux observed in the microscopy images of CTL and epiCTL are reflected in the cellular media, a sample of 100 µL was taken before adding the media to the cells (start), 24 hrs after uptake media was added (uptake), and after 24 hrs of efflux (efflux). Similar samples were taken from cells treated with only BSA without loading of CTL and epiCTL, and these spectra were subtracted from the ones of CTL and epiCTL. The 100 µL samples were diluted with 400 µL of 1X PBS before they were added to the cuvette in the spectrofluorometer. Fig. S2A and B show the spectra recorded for CTL and epiCTL, respectively. Fig. S2C shows the integrated intensity of each spectra, and Fig. S2D shows the percentage uptake and efflux for CTL and epiCTL. The percentage uptake is calculated as:

$$\text{Percentage uptake} = \frac{\sum \text{Start} - \sum \text{Uptake}}{\sum \text{Start}} \cdot 100 \quad \textbf{(1)}$$

Since $\sum$Start is the integrated intensity of the fluorescent sterol before any uptake, and $\sum$Uptake is the remaining sterol left in the media after uptake, the difference gives the actual uptake of the sterol. The percentage effluxed is then calculated by:

$$\text{Percentage efflux} = \frac{\sum \text{Efflux}}{\sum \text{Start} - \sum \text{Uptake}} \cdot 100 \quad \textbf{(2)}$$

This shows a similar uptake of CTL and epiCTL, but a lower efflux of epiCTL.

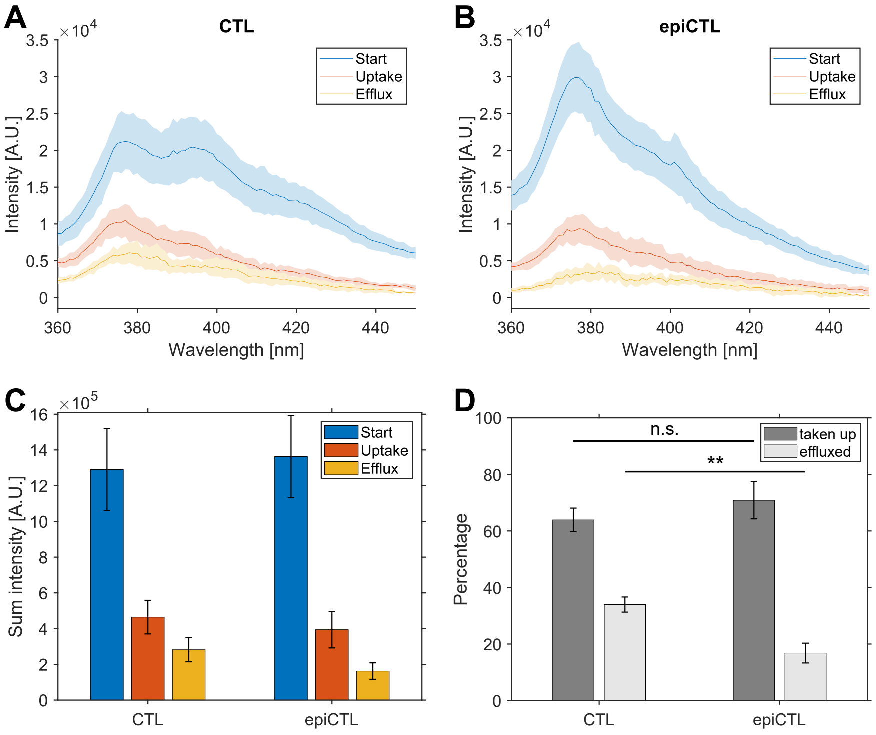


**Fig. S2.** (A and B) Fluorescence spectra of CTL and epiCTL acquired before uptake, after uptake, and after efflux, respectively. The spectra are shown as the mean of three measurements, and the shaded area around the curve represents the SD. Excitation at 335 nm, emission measured from 360-450 nm with a step size of 1 nm, and an integration time of 1 s. (C) Bar chart showing the intensity of the integrated spectra shown in **A** and **B**. (D) Bar chart showing the percentage uptake and efflux for CTL and epiCTL. The bars show a mean of three with error bars depicting the SD. A two-sample t-test shows no difference between the uptake of CTL and epiCTL (p-value: 0.1976), but a ** difference between the efflux percentage of CTL and epiCTL (p-value: 0.0025).

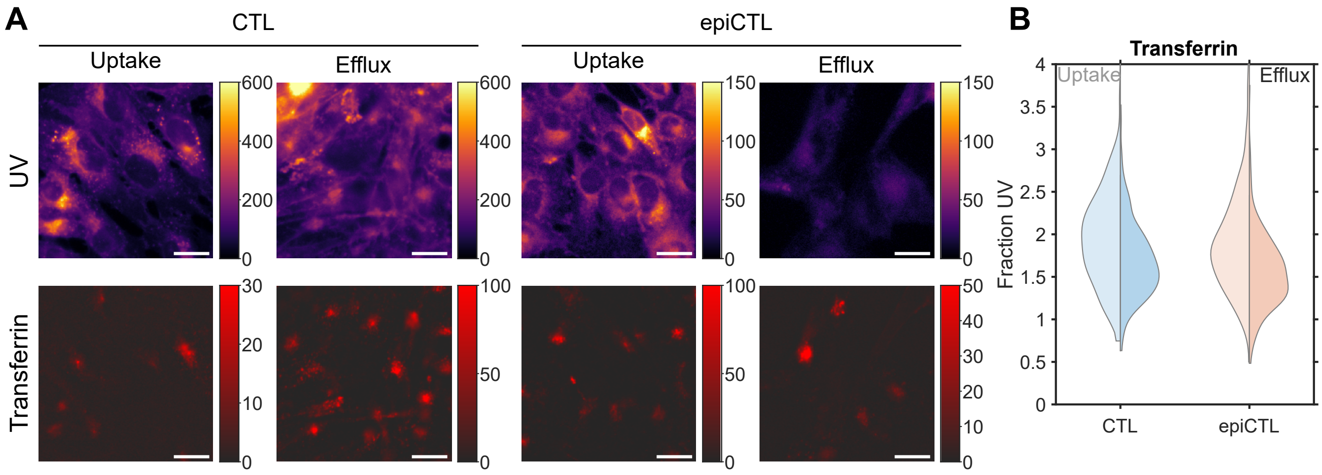


**Fig. S3.** (A) Fluorescence widefield images of CTL or epiCTL in the top row taken after uptake and efflux, and transferrin-CF640R in the bottom row for the same images (Scale bars 20 µm). (B) Double violin plot showing the UV fraction of either CTL or epiCTL in the transferrin channel.

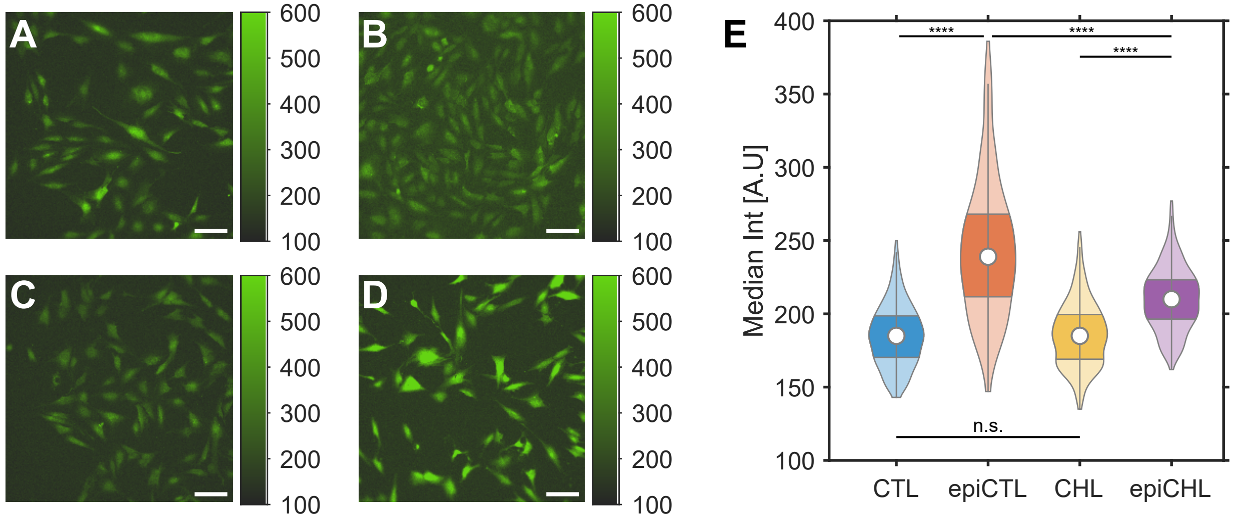


**Fig. S4.** Higher cytosolic calicium in cells treated with epiCTL. Representative fluorescence widefield images of IM-NHA labeled with 0.5 $\mu$M of Cal520-AM after 24 hr treatment with either (A) CHL/BSA, (B) epiCHL/BSA, (C) CTL/BSA, or (D) epiCTL/BSA for 24 hrs. Scale bars are 100 µm. E) Violin-plot showing the distributions of median cell intensity. Cells are pooled together from 2 experiments, CTL $N_{cells}$ = 1020, epiCTL $N_{cells}$ = 1028, CHL $N_{cells}$ = 1049, and epiCHL $N_{cells}$ = 1149.

**Distribution of CTL and epiCTL under efflux conditions.**

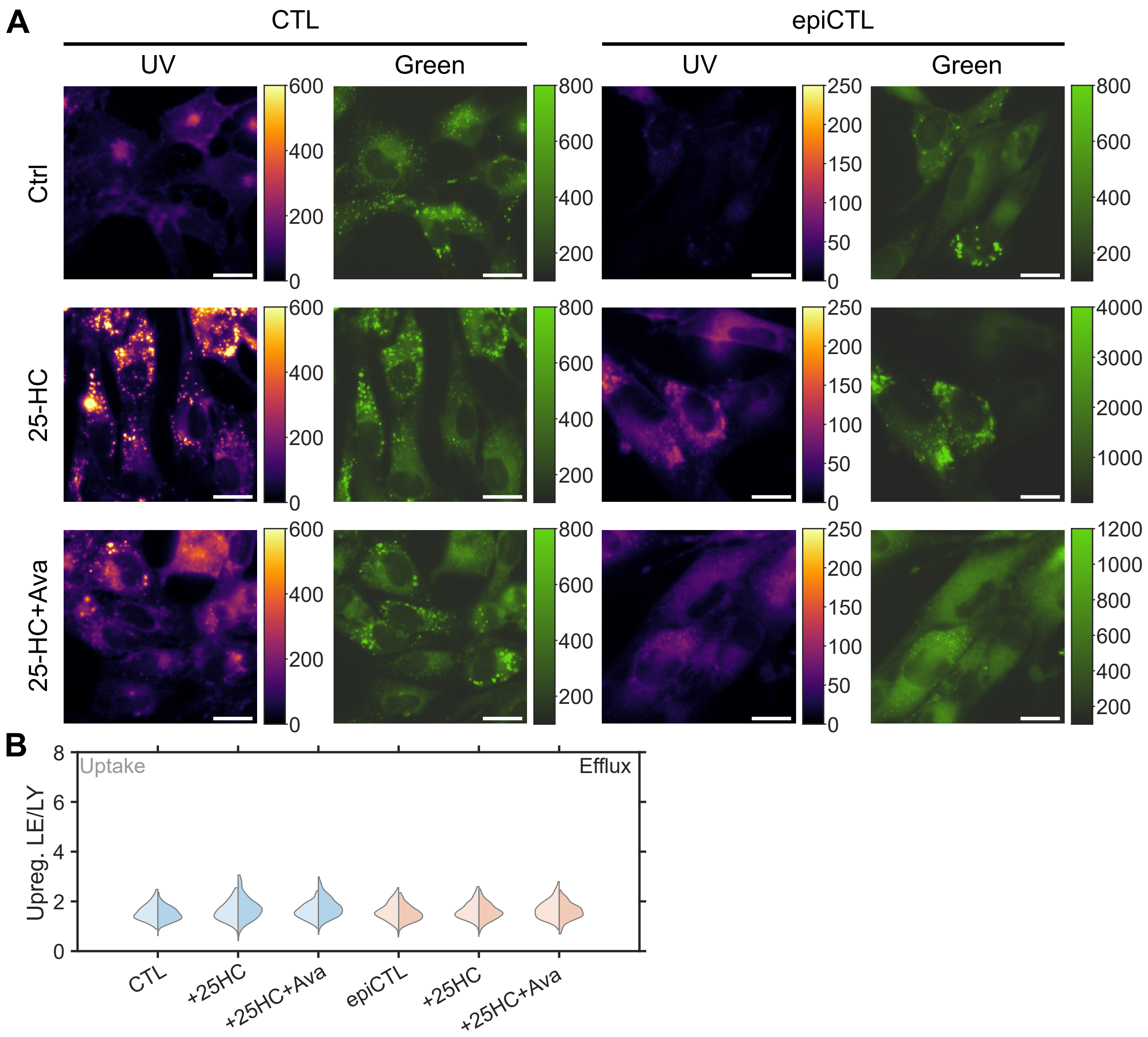


**Fig. S5.** 25HC reveals stereospecific esterification of CTL, but not epiCTL during efflux, without affecting LE/LY distribution. (A) WF fluorescence images were taken in the UV channel (CTL), green channel (Bodipy), and red channel (RhDex) of IM-NHA loaded with CTL/BSA for 24 hrs (uptake) and allowed to efflux for 24 hrs. Scale bars are 20 µm. (B) Double-sided violin plot showing the mean fraction of the UV signal found in the LE/LY in uptake (left) versus efflux (right) for CTL and epiCTL.

**Transport of DHE by SBDs.** Interestingly, transport of DHE (for structure see Fig. S6A) by the investigated SBDs is also different to what is observed for CTL and epiCTL. Here, Aster-B proves to be the most efficient, then Aster-A, and last Aster-C (Fig. S6B). For the STARD-proteins STARTD1 and STARD4 prove to be very efficient for DHE (Fig. S6C), whereas STARD 5 alsmost does not transfer DHE but is very efficient for both CTL and epiCTL (Fig. 3D and E).

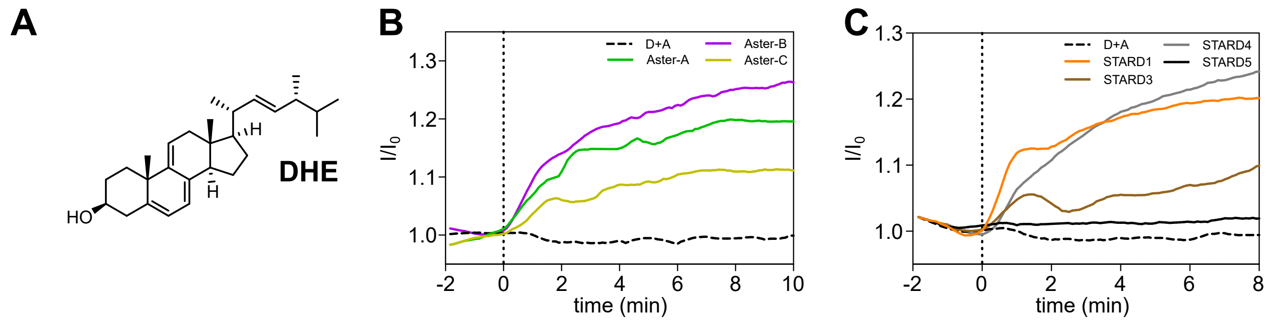


**Fig. S6.** Characterization of transport of DHE by SBDs. (A) Structure of DHE. (B) Transport of DHE by 1 µM Aster-A/B/C, as assessed by a FRET assay. (C) Transport of DHE by 1 µM STARD1/3/4/5, as assessed by a FRET assay. All data represent a representative experiment from two biological replicates (n = 2).

| Protein | DHE | CTL | epiCTL |
|---|---|---|---|
| Aster A | ++ | +++ | ++ |
| Aster B | +++ | ++ | + |
| Aster C | + | + | ++ |
| StARD1 | +++ | +++ | - |
| StARD3 | + | - | - |
| StARD4 | +++ | + | ++ |
| StARD5 | - | +++ | +++ |

**Table 1.** Overview of transport ability of the different proteins to DHE, CTL, and epiCTL. Where '+++' is being very efficient, '++' is medium efficient, '+' can transfer, and '-' no transfer.
